# Dual roles of EGO-1 and RRF-1 in regulating germline exo-RNAi efficiency in *Caenorhabditis elegans*

**DOI:** 10.1101/2024.09.24.614643

**Authors:** Katsufumi Dejima, Keita Yoshida, Shohei Mitani

## Abstract

RNA interference (RNAi) is widely used in life science research and is critical for diverse biological processes, such as germline development and antiviral defense. In *Caenorhabditis elegans*, RNA-dependent RNA polymerases, with redundant involvement of EGO-1 and RRF-1, facilitate small RNA amplification in germline exogenous RNAi (exo-RNAi). However, their coordination during the regulation of exo-RNAi processes in the germline remains unclear. Here, we examined non-null mutants of the *ego-1* gene and found that *ego-1(S1198L)* animals exhibited germline exo-RNAi defects with normal fertility, abnormalities in germ granules, and synthetic temperature-dependent sterility with *rrf-1*. The exo-RNAi defects in *ego-1(S1198L)* were partially restored by inhibiting *hrde-1*, *cde-1*, and *znfx-1*. Similar defects were observed in wild-type and *ego-1(S1198L)* heterozygous descendants derived from *ego-1(S1198L)*, but these were suppressed by ancestral inhibition of *rrf-1*. These data reveal a dual role for EGO-1 in the positive regulation of germline exo-RNAi: it not only mediates target silencing through its RNA-dependent RNA polymerase activity, but also fine-tunes germ granule function or downstream processes, which are antagonized by RRF-1.

## Introduction

RNA interference (RNAi) is a process in which double-stranded RNA (dsRNA) triggers the silencing of mRNAs with sequence homology to the dsRNA (Fire *et al*, 1998). RNAi plays a role in the regulation of gene expression, RNA quality control, and anti-transposon and antiviral responses in many eukaryotes, as well as in epigenetic inheritance in certain eukaryotes (Castel & Martienssen, 2013). RNAi can be induced using endogenous or exogenous dsRNA sources. Regardless of whether initiated by an exogenous or endogenous source, RNAi in *Caenorhabditis elegans* is a common process that leads to the amplification of secondary small RNAs (sRNAs) by RNA-dependent RNA polymerases (RdRPs). In exogenous RNAi (exo-RNAi), the endonuclease dicer cleaves the exogenously introduced long dsRNAs into primary sRNAs (Tabara *et al*, 2002). The primary Argonaute protein RDE-1 loads the resulting sRNAs and recruits RdRPs to produce secondary sRNA populations that regulate their target mRNA (Sijen *et al*, 2001; Tabara *et al*, 1999). In endogenous RNAi, several primary Argonaute proteins, including CSR-1, PRG-1, ERGO-1, and ALG-3/4, selectively bind endogenous non-coding RNAs such as 21U-RNA and 26G RNA (Batista *et al*, 2008; Claycomb *et al*, 2009; Conine *et al*, 2010; Das *et al*, 2008; Gent *et al*, 2010; Pavelec *et al*, 2009; Wang & Reinke, 2008). The 21U-RNA is transcribed from the 21U-RNA locus by RNA polymerase II under the regulation of FKH transcription factors, whereas 26G RNA is a type of secondary RNA synthesized by the <u>e</u>nhanced <u>R</u>NAi (ERI) complex using spliced mRNA as a template (Cecere *et al*, 2012; Kennedy *et al*, 2004). The binding process between primary Argonaute proteins and these sRNAs facilitates targeting of the corresponding RNAs. This interaction recruits the RdRP protein, leading to the amplification of secondary sRNAs. There are four RdRPs in *C. elegans*, including EGO-1, RRF-1, RRF-2, and RRF-3 (Sijen *et al*., 2001; Smardon *et al*, 2000), all of which are involved in sRNA synthesis in the endogenous RNAi pathway. RRF-2 and RRF-3 are involved in the amplification of risi-RNA and 26G-sRNA, respectively (Simmer *et al*, 2002; Zhou *et al*, 2017). EGO-1 mediates both CSR-1 and WAGO 22G RNA amplification, whereas RRF-1 mediates WAGO 22G RNA and risi-RNA amplification (Claycomb *et al*., 2009; Gu *et al*, 2009; Zhou *et al*., 2017).

In germline exo-RNAi, an intrinsic amplification system mediated by EGO-1 and RRF-1 enables efficient gene knockdown. RRF-1 functions in both germline and somatic cells, whereas EGO-1 functions exclusively in germline cells, with partial involvement in some somatic cells (Kumsta & Hansen, 2012). The *ego-1* and *rrf-1* genes are physically close to the genome and are expected to be transcribed together as operons. Thus, they may exhibit similar epigenetic and transcriptional regulations. However, it is likely that their functional differences are not solely due to transcriptional regulation but rather arise from the specific roles played by each molecule. While RRF-1 has the ability to synthesize mutator complex-dependent sRNA, which is critical for gene silencing in exo-RNAi, EGO-1 plays a compensatory role in promoting germline exo-RNAi, complementing the function of RRF-1 (Claycomb *et al*., 2009; Gu *et al*., 2009; Zhou *et al*., 2017). This idea is supported by two key findings: (i) *rrf-1* null mutants are exo-RNAi sensitive in the germline (Sijen *et al*., 2001); and (ii) in the absence of RRF-1, EGO-1 is recruited to the mutator complex to compensate for RRF-1’s function (Phillips *et al*, 2012; Phillips & Updike, 2022). However, *ego-1* null mutants exhibit exo-RNAi germline defects, whereas *rrf-1* mutants do not, and the exact reason for this difference remains unclear. Null mutations in *ego-1* result in severe germline developmental abnormalities, making it challenging to conduct detailed analyses, including assessment of exo-RNAi effects.

Understanding the role of germ granules is crucial for studying germline exo-RNAi. P granules safeguard the transcripts of germline-expressing genes against silencing mediated by PRG-1 and HRDE-1/WAGO-9 (Dodson & Kennedy, 2019; Ouyang *et al*, 2019). This protective mechanism is important in exo-RNAi activity, as the safeguarded transcripts include the RNAi genes *sid-1* and *rde-11* (Dodson & Kennedy, 2019; Lev *et al*, 2019; Ouyang *et al*., 2019). Transcripts of *sid-1* and *rde-11* accumulate in P granules, and the loss of P granules results in their dispersion and aberrant levels of WAGO 22G RNA associated with HRDE-1 (Dodson & Kennedy, 2019; Ouyang *et al*., 2019). This, in turn, results in the silencing of *sid-1* and *rde-11* genes and defects in germline exo-RNAi. In contrast, CSR-1 opposes PRG-1/piRNA complex activity, which triggers HRDE-1-mediated transgenerational gene silencing (Shen *et al*, 2018). Therefore, as both WAGO and CSR-1 class sRNAs are involved, EGO-1 and RRF-1 seem to have differential roles in synthesizing the sRNAs loaded onto these Argonauts. Recently, inhibition of E granule biogenesis, which includes EGO-1, has been shown to upregulate sRNA targeting *sid-1* and *rde-11*, leading to silencing of these genes (Chen *et al*, 2024). However, the orchestration of these complex regulatory mechanisms in germline exo-RNAi remains unknown.

Here, we examined non-null alleles for the *ego-1* gene from the million mutation project (MMP) collection (Thompson *et al*, 2013). Four alleles showed germline exo-RNAi defects with normal fertility at 20°C, in contrast to *ego-1* null animals. We further analyzed the mutant strain with *ego-1(*S1198L) and found synthetic effects on the temperature-sensitive sterile phenotype of *ego-1(*S1198L) by *rrf-1* deletion mutation. The levels of GFP::ZNFX-1, PGL-1::mCherry, and HRDE-1::GFP increased in *ego-1(*S1198L) pachytene stage cells. In addition, we found that the transcripts of *sid-1* and *rde-11* in *ego-1(*S1198L) were downregulated, which was suppressed in *hrde-1*, *cde-1*, and *znfx-1* mutants. Consistently, the RNAi-defective (Rde) phenotype detached from the *ego-1(*S1198L) genotype over generations in an RRF-1 dependent manner. Our data demonstrate that EGO-1 plays a role in enhancing the robustness of exo-RNAi in the germline by mediating at least two processes: it acts as an RdRP necessary for target gene silencing by complementing RRF-1, and it facilitates the regulation of exo-RNAi gene expression in the germline. These findings reveal that an extensive interdependent RdRP network is responsible for regulating germline exo-RNAi.

## Results

### Some non-null *ego-1* alleles show germline exo-RNAi defects

Null mutants of the RdRP gene, *ego-1*, exhibit a completely sterile phenotype (Smardon *et al*., 2000). Within the MMP strain collection, 23 viable and fertile strains have been identified, each carrying a missense mutation at the *ego-1* locus (Table 1) (Dejima & Mitani, 2022; Thompson *et al*., 2013). Single amino acid substitutions were distributed throughout the EGO-1 protein, including the RdRP and coiled-coil domains (Fig. 1A). Bioinformatic programs commonly used to predict the effects of missense mutations suggested that some mutations were likely to have deleterious effects (Table 1) (Adzhubei *et al*, 2010; Choi *et al*, 2012; Ng & Henikoff, 2001). Given the critical role of the EGO-1 protein in RNAi, we investigated whether germline exo-RNAi was defective in all mutants. To investigate this phenotype, we fed these *ego-1* mutant hermaphrodites with bacteria expressing dsRNA against *pos-1* or *pop-1*. The *pos-1* and *pop-1* genes are maternally expressed in germ cells and are essential for embryonic viability. Among the mutant strains that were outcrossed with the fluorescent chromosomal balancer *tmC18[tmIs1200]* (Dejima & Mitani, 2022), *ego-1(gk357146)*, *ego-1(gk426642)*, *ego-1(gk532049),* and *ego-1(gk882383)* showed an Rde phenotype when they were fed with the HT115 expressing germline gene, either *pos-1* or *pop-1* dsRNA (Table 1 and Fig. 1B, C). Hereafter, we refer to *ego-1(gk357146)* as *ego-1*(V1128E), *ego-1(gk426642)* as *ego-1*(R539Q), *ego-1(gk532049)* as *ego-1(S1198L)*, and *ego-1(gk882383)* as *ego-1*(C823Y).

**Figure 1.**
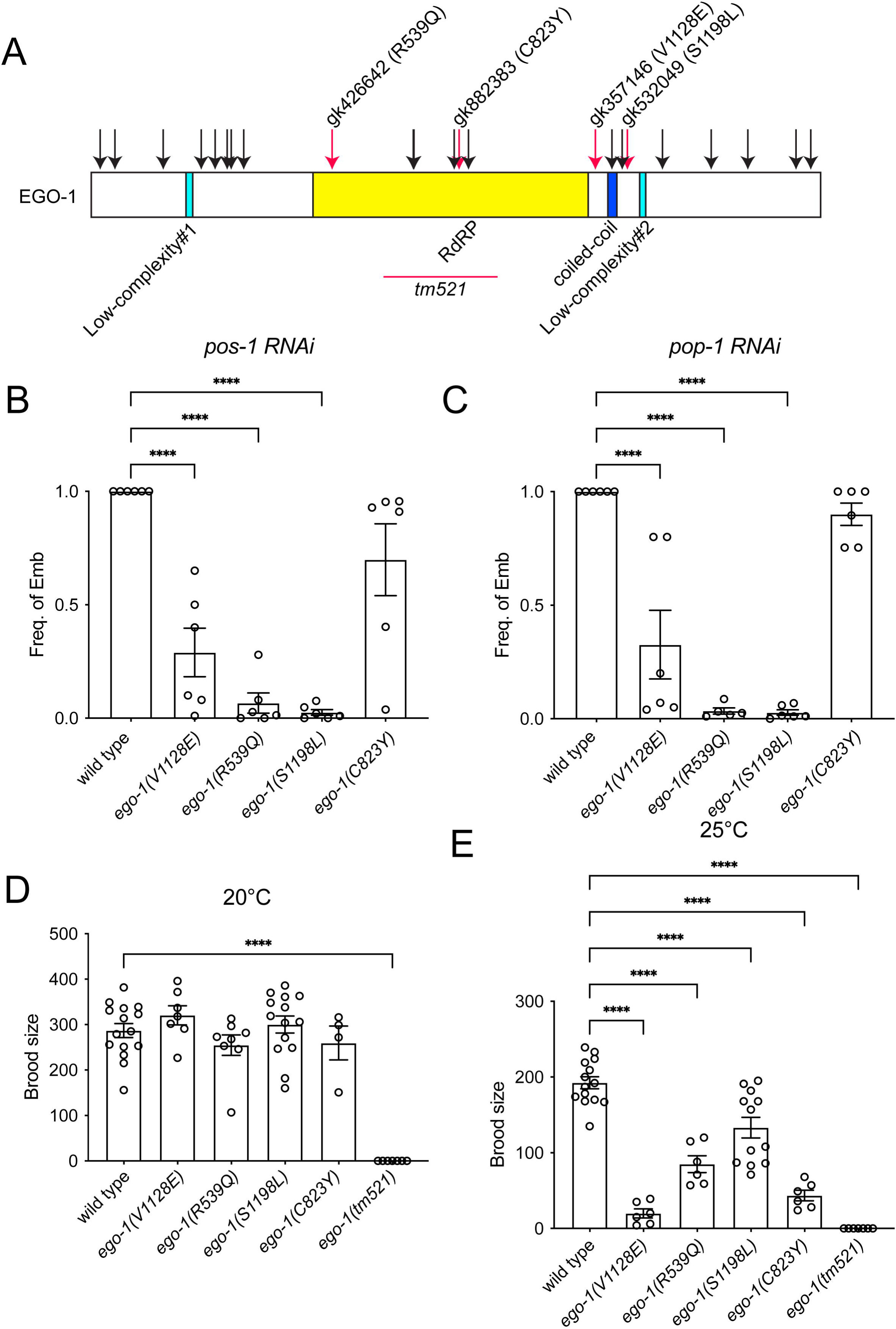
The non-null mutations of *ego-1*, which cause germline exo-RNAi defects. A. Protein structure of EGO-1 showing mutations that were used in this study. The mutations that cause germline exo-RNAi defects are indicated by red arrows, while those that do not cause such defects are marked with black arrows. B. Quantification of the frequency of indicated mutant animals showing the Emb phenotype under *pos-1* feeding RNAi. C. Quantification of the frequency of indicated mutant animals showing the Emb phenotype under *pop-1* feeding RNAi. D and E. Brood size analysis of the indicated mutant animals that were cultured at 20°C (D) and 25°C (E).

**Table 1.**

| Allele name | Amino acid change | Amino acid aligned in RRF-1 | PolyPhen-2 | SHIFT | PROVEAN | *pos-1* RNAi (if not Rde, %Emb ± SD is shown) | *pop-1* RNAi (if not Rde, %Emb ± SD is shown) |
| --- | --- | --- | --- | --- | --- | --- | --- |
| *gk115401* | E20K | S | benign | NA | neutral | 100±0 | 100±0 |
| *gk720210* | V53I | I | benign | NA | neutral | 100±0 | 100±0 |
| *gk470185* | P161S | - | benign | NA | neutral | 100±0 | 100±0 |
| *gk498425* | L246F | V | probably damaging | NA | Deleterious | 98.3±1.5 | 100±0 |
| *gk115397* | V277M | C | possibly damaging | NA | neutral | 100±0 | 100±0 |
| *gk115395* | D304N | H | benign | NA | neutral | 100±0 | 100±0 |
| *gk837385* | Y313F | Y (conserved) | probably damaging | NA | neutral | 100±0 | 100±0 |
| *gk115394* | R341K | S | benign | NA | neutral | 100±0 | 100±0 |
| *gk426642* | R539Q | R | possibly | TOLERATED | Deleterious | Rde (data is | Rde (data is |
|  |  | (conserved) | damaging |  |  | shown in<br>Fig. 1B and<br>C) | shown in<br>Fig. 1B and<br>C) |
| *gk896494* | L721F | I | benign | TOLERATED | neutral | 96.7±5.8 | 100±0 |
| *gk115393* | R722H | R<br>(conserved) | benign | DELETERIOUS | neutral | 100±0 | 100±0 |
| *gk481348* | S812F | S<br>(conserved) | benign | DELETERIOUS | Deleterious | 100±0 | 100±0 |
| *gk882383* | C823Y | C<br>(conserved) | probably<br>damaging | TOLERATED | Deleterious | Rde (data is<br>shown in<br>Fig. 1B and<br>C) | Rde (data is<br>shown in<br>Fig. 1B and<br>C) |
| *gk721963* | G844E | G<br>(conserved) | probably<br>damaging | DELETERIOUS | Deleterious | 100±0 | 100±0 |
| *gk357146* | V1128E | V<br>(conserved) | possibly<br>damaging | TOLERATED | Deleterious | Rde (data is<br>shown in<br>Fig. 1B and<br>C) | Rde (data is<br>shown in<br>Fig. 1B and<br>C) |
| *gk925207* | R1165K | S | benign | TOLERATED | neutral | 100±0 | 100±0 |
| *gk749674* | G1188E | G<br>(conserved) | probably<br>damaging | DELETERIOUS | Deleterious | 98±2 | 96.3±3.2 |
| *gk532049* | S1198L | S<br>(conserved) | possibly<br>damaging | TOLERATED | Deleterious | Rde (data is<br>shown in<br>Fig. 1B and<br>C) | Rde (data is<br>shown in<br>Fig. 1B and<br>C) |
| *gk115391* | E1278K | E<br>(conserved) | benign | TOLERATED | neutral | 100±0 | 100±0 |
| *gk317493* | E1397K | I | benign | NA | neutral | 99.7±0.6 | 100±0 |
| *gk115390* | K1469E | K<br>(conserved) | possibly<br>damaging | NA | neutral | 100±0 | 100±0 |
| *gk864727* | N1577S | T | benign | NA | neutral | 100±0 | 100±0 |
| *gk540555* | G1610R | G<br>(conserved) | probably<br>damaging | NA | neutral | 100±0 | 100±0 |

### Mutants with the exo-RNAi defective *ego-1* allele are fertile but show a temperature-sensitive sterile phenotype in the absence of RRF-1

All of these germline exo-RNAi defective mutants showed normal brood size at 20°C, whereas the *ego-1(tm521)* deletion mutant showed a completely sterile phenotype (Fig. 1D). The temperature-sensitive sterile phenotype is a hallmark of defects in the mutator complex genes required for WAGO 22G RNA production (Ketting *et al*, 1999), which is necessary for exo-RNAi-driven silencing. To evaluate whether the exo-RNAi-defective phenotype in these *ego-1* mutants was due to defects in WAGO 22G RNA production, we examined the brood size of these mutants at high temperatures (Fig. 1E). The *ego-1* mutants that exhibited the strong germline exo-RNAi defective phenotype (R539Q and S1198L) showed mild defects in brood size at 25°C, suggesting that the production of WAGO 22G RNA produced by the mutator complex was not completely disrupted in these mutants. Since RRF-1 was shown to be primarily included in the mutator complex, we investigated whether the loss of *rrf-1* affected fertility in *ego-1*(S1198L). For further analysis, we focused on *ego-1*(S1198L), which exhibited the most pronounced Rde phenotype. We knocked out the *rrf-1* gene, which is adjacent to the *ego-1* gene in terms of its genomic locus, in the *ego-1*(S1198L) background using the CRISPR/Cas9 system to create *rrf-1(tm9941) ego-1(*S1198L) double mutants (Fig. 2A). For comparison, we generated a similar deletion allele, *rrf-1(tm9951)* (Fig. 2A). While *ego-1*(S1198L) was sensitive to somatic RNAi, *rrf-1(tm9951)* and *rrf-1(tm9941) ego-1(*S1198L) mutants were completely defective in somatic RNAi (Fig. EV1A, B). The *rrf-1* single deletion mutant *rrf-1(tm9951)*, as well as a commonly used and larger deletion allele *rrf-1(pk1417)* (Sijen *et al*., 2001), showed normal brood size at both 20°C and 25°C (Fig. 2B, C). In contrast, *rrf-1(tm9941) ego-1(*S1198L) double mutant homozygotes showed temperature-sensitive sterility similar to *mut-14(pk738)* and *mut-15(tm1358)* mutants (Fig. 2B, C), suggesting that thermotolerant gametogenesis in *ego-1(*S1198L) is maintained by a redundant function of RRF-1 in an endogenous RNAi mechanism mediated by the mutator complex.

**Figure 2.**
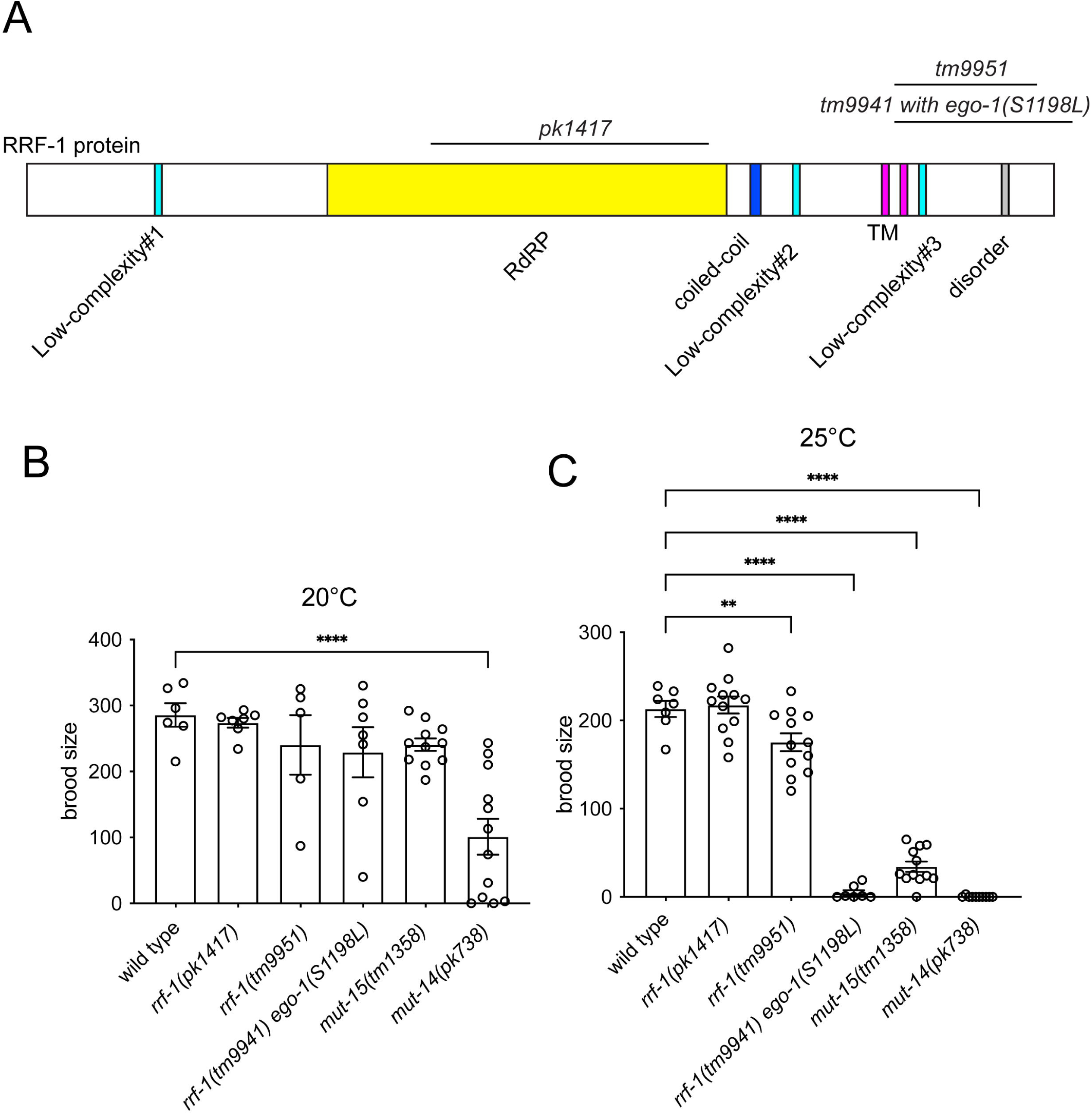
Combining the *ego-1(S1198L)* mutation with the *rrf-1* deletion mutation results in a temperature-sensitive sterile phenotype. A. Protein structure of RRF-1 showing the mutations used in this study. TM: transmembrane domain. B and C. Brood size analysis of the indicated mutant animals that were cultured at 20°C (B) and 25°C (C).

### Increased intensity of *ego-1(S1198L)* germ granules in pachytene-stage cells

Previous studies have implicated EGO-1 in P granule biogenesis (Vought *et al*, 2005). P granules serve as safe harbors where germline-expressed mRNA is protected from silencing by PRG-1 and HRDE-1 (Dodson & Kennedy, 2019) (Ouyang *et al*., 2019) (Lev *et al*., 2019). Additionally, it has been shown that EGO-1 binds to ZNFX-1, a Z granule component required for RNAi inheritance, together with HRDE-1 (Ishidate *et al*, 2018; Ouyang *et al*, 2022). To determine whether these granules were affected, we analyzed the patterns of PGL-1::tagRFP and GFP::ZNFX-1 in the germline. Perinuclear ZNFX-1 and PGL-1 puncta were observed in the wild-type pachytene region. The overall pattern of these puncta was not affected in *ego-1(S1198L)* homozygotes (Fig. 3A, B). However, in *ego-1(S1198L)* homozygotes, the fluorescence intensity per punctum significantly increased for both PGL-1::tagRFP and GFP::ZNFX-1 (Fig. 3C, D).

**Figure 3.**
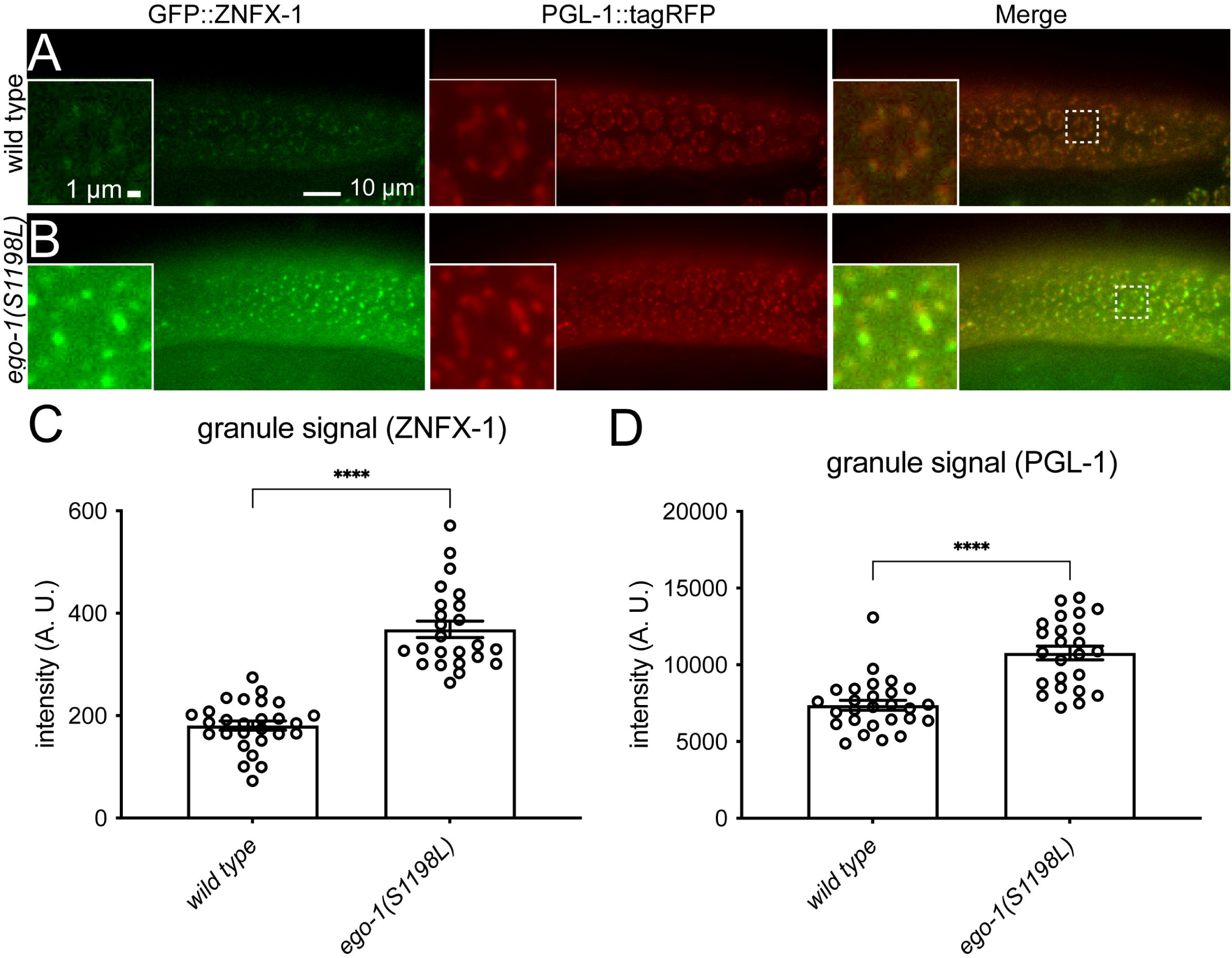
Expression of GFP::ZNFX-1 and PGL-1::tagRFP in *ego-1(S1198L).* A and B. Fluorescence images of germ cells showing the localization of GFP::ZNFX-1 and PGL-1::tagRFP in the wild-type (A) and *ego-1(S1198L)* (B). C and D. Bar graphs show the quantification of granular GFP::ZNFX-1 (C) and PGL-1::tagRFP (D) fluorescence signal intensity.

### Loss of CDE-1 or ZNFX-1 partially suppresses the germline exo-RNAi defects of *ego-1(S1198L)*

Since the fluorescence intensity per punctum for GFP::ZNFX-1 increased in *ego-1(S1198L)* homozygotes, we next examined germline exo-RNAi activity in double mutants for *ego-1(S1198L)* and *znfx-1(gg561)*. *znfx-1(gg561)* suppressed the Rde phenotype of *pos-1* RNAi in *ego-1(S1198L)* (Fig. 4A). Although not statistically significant, *znfx-1(gg561)* slightly increased sensitivity to *pop-1* RNAi (Fig. 4B). Therefore, a plausible interpretation is that the *ego-1(S1198L)* mutation upregulates the functionality of ZNFX-1 and/or Z granules.

**Figure 4.**
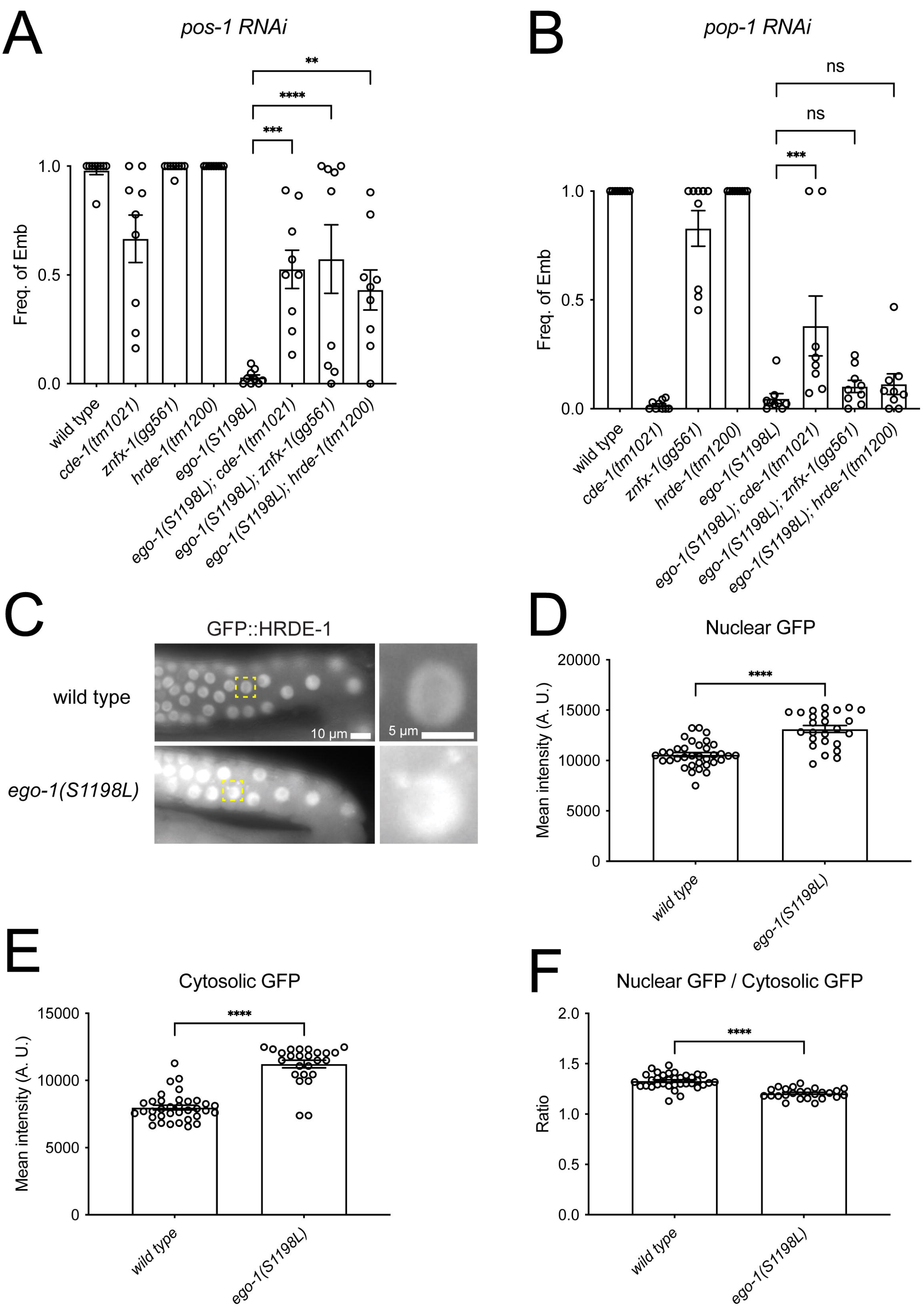
Germline exo-RNAi defects in *ego-1(S1198L)* are partially suppressed by *cde-1* and *hrde-1* mutations. A. Quantification of the frequency of indicated mutant animals showing the Emb phenotype under *pos-1* feeding RNAi. B. Quantification of the frequency of indicated mutant animals showing the Emb phenotype under *pop-1* feeding RNAi. C. Fluorescence images of germ cells showing the localization of GFP::HRDE-1. D. Bar graph showing the quantification of nuclear GFP::HRDE-1 fluorescence signal intensity. E. Bar graph showing the quantification of cytoplasmic GFP::HRDE-1 fluorescence signal intensity. F. Bar graph showing the ratio of nuclear GFP/cytoplasmic GFP signals.

Based on the fact that ZNFX-1 physically interacts with EGO-1(Ishidate *et al*., 2018), we focused on other molecules that interact with EGO-1. Previous studies have shown that EGO-1 directly interacts with the germline-specific polyuridylation polymerase CDE-1/CID-1 (van Wolfswinkel *et al*, 2009; Xu *et al*, 2018). Although, for unknown reasons, *cde-1(tm1021)* displayed a full Rde phenotype in response to *pop-1* RNAi, it was sensitive to *pos-1* RNAi (Fig. 4A, B). Importantly, *cde-1(tm1021)* partially suppressed the Rde phenotype in response to both *pos-1* and *pop-1* RNAi in *ego-1(*S1198L) (Fig. 4B). Uridylation by CDE-1 stabilizes WAGO 22G and destabilizes CSR-1 22G RNAs (van Wolfswinkel *et al*., 2009) (Xu *et al*., 2018). Loss of CDE-1 increased 22G RNA loading on CSR-1 and decreased 22G RNA loading on WAGOs other than CSR-1. Thus, *ego-1(S1198L)* could potentially lead to an imbalance in the loading of 22G RNA onto these Argonauts, in contrast to the effect of CDE-1 loss.

### Loss of HRDE-1 partially suppresses the germline exo-RNAi defects in *ego-1(S1198L)*

HRDE-1 is a nuclear Argonaute protein that mediates nuclear RNAi and inheritance of piRNA silencing in germ cells (Ashe *et al*, 2012; Buckley *et al*, 2012; Luteijn *et al*, 2012; Shirayama *et al*, 2012). HRDE-1-dependent gene silencing has been shown to inhibit the RNA polymerase II-dependent RNA transcription of germline genes, including *sid-1* and *rde-11*, which are required for exo-RNAi (Dodson & Kennedy, 2019; Ouyang *et al*., 2019). Suppression of the Rde phenotype, which was observed in *ego-1(S1198L); cde-1* double mutants, suggests that the loading of 22G RNA onto WAGOs, including HRDE-1, other than CSR-1 may be increased in *ego-1(S1198L)*. To test whether the Rde phenotype was caused by the HRDE-1-mediated silencing of germline exo-RNAi genes, we examined the genetic interactions between *hrde-1(tm1200)* and *ego-1(S1198L)*. The *hrde-1* single mutants showed normal responses to *pos-1* and *pop-1* RNAi, as reported previously (Yigit *et al*, 2006). Similar to *cde-1(tm1021)* and *znfx-1(gg561)*, *hrde-1(tm1200)* partially suppressed the Rde phenotype of *pos-1* RNAi in *ego-1(*S1198L) (Fig. 4A).

### HRDE-1::GFP accumulates in *ego-1(*S1198L) pachytene-stage cells

Since germline exo-RNAi defects in *ego-1(S1198L)* were suppressed in the absence of HRDE-1, we wondered whether the distribution of the HRDE-1 protein was altered in *ego-1(S1198L)*. HRDE-1::GFP was predominantly localized to the nuclei in wild-type pachytene-stage cells, as reported previously (Fig. 4C) (Ashe *et al*., 2012; Buckley *et al*., 2012). In *ego-1(S1198L)* homozygous pachytene-stage cells, both nuclear and cytosolic levels of HRDE-1::GFP increased, and germ granule-like dots were detected (Fig. 4C, D). Additionally, the nuclear/cytosolic ratio decreased slightly (Fig. 4E). These observations suggest that EGO-1, by predominantly producing CSR-1 22G RNA, possibly limits the loading of 22G RNA onto HRDE-1, which, in turn, affects the proper subcellular distribution of HRDE-1.

### The expression of *sid-1* and *rde-11* is downregulated in *ego-1(S1198L)* in an HRDE-1-dependent manner

To evaluate the expression of *sid-1* and *rde-11* in the mutants, RT-qPCR was performed. Expression of these genes was significantly downregulated in the *ego-1(*S1198L) mutant (Fig. 5A, B). Similarly, expression of these genes was significantly downregulated in the *ego-1(tm521)* mutant (Fig. 5C, D). The reduced levels observed in *ego-1(*S1198L) and *ego-1(tm521)* were comparable, suggesting that the *ego-1(S1198L)* mutation induced defects equivalent to loss of function in terms of the regulation of gene expression. The expression of these genes in the double mutants for *ego-1(*S1198L) and *hrde-1(tm1200)* and *ego-1(*S1198L) and *cde-1(tm1021)* was restored to wild-type levels (Fig. 5A, B). These data suggest that EGO-1 affects the regulation of 22G RNA loading on HRDE-1, which in turn contributes to the germline expression of *sid-1* and *rde-11*.

**Figure 5.**
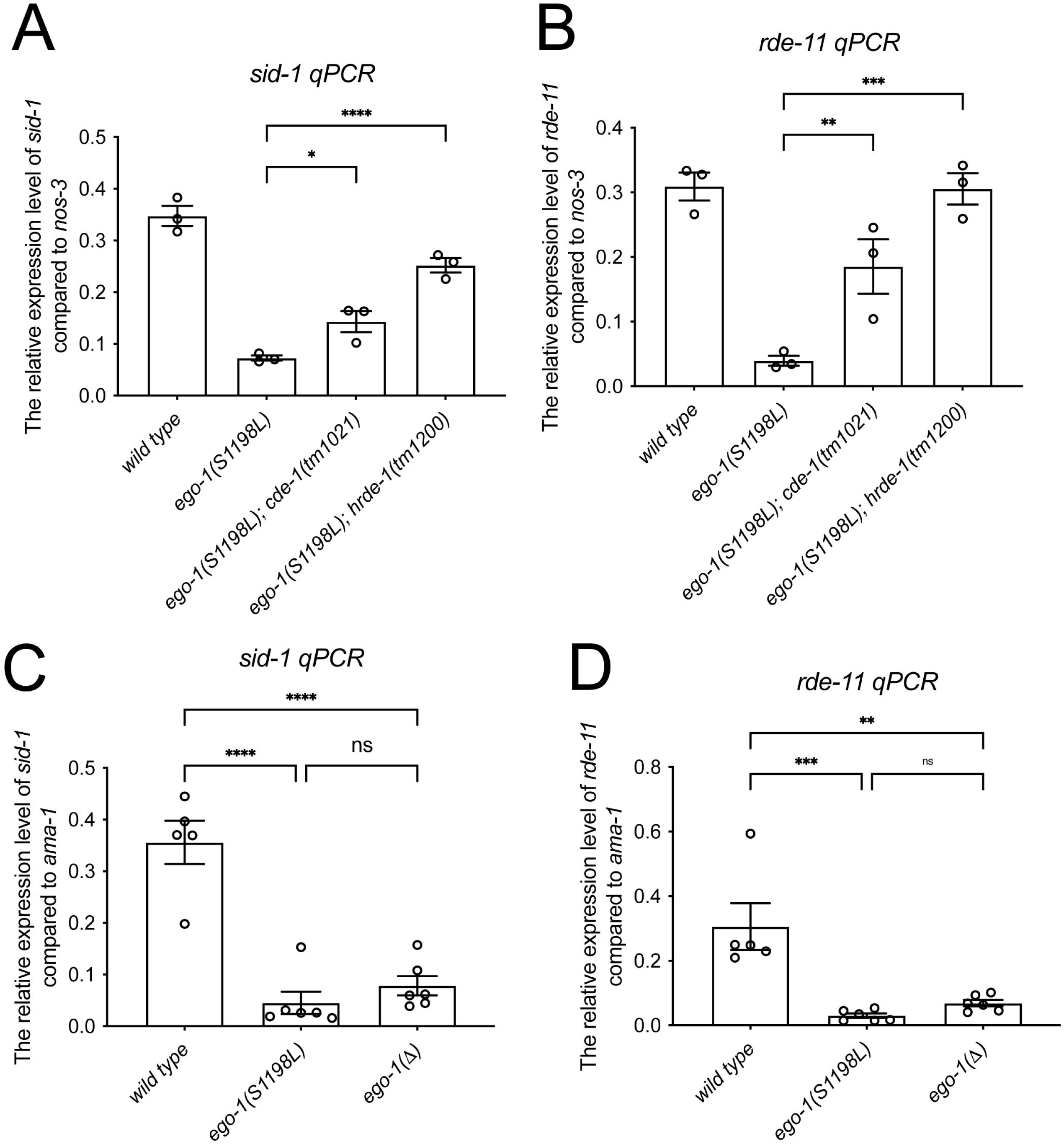
*ego-1(S1198L)* shows reduced expression of *sid-1* and *rde-11* in a CDE-1- and HRDE-1-dependent manner. A and B. RT-qPCR showing the relative expression levels of *sid-1* (A) and *rde-11* (B) mRNA normalized to *nos-3* expression. RNA was extracted from young-adult animals grown at 20°C. C and D. RT-qPCR showing the relative expression levels of *sid-1* (E) and *rde-11* (F) mRNA normalized to *ama-1* expression. RNA was extracted from young-adult animals grown at 20°C.

### Characterization of germline exo-RNAi phenotypes in *ego-1* mutant heterozygotes

Since we observed compensation by RRF-1 in temperature-sensitive sterility, but not in germline exo-RNAi, the defects of *ego-1(*S1198L) in germline exo-RNAi could manifest as inhibition of an exo-RNAi process that requires both EGO-1 and RRF-1. In this scenario, *ego-1(*S1198L) should be genetically dominant among germline exo-RNAi. To address this, we examined the germline exo-RNAi sensitivity of *ego-1(*S1198L) heterozygous hermaphrodites. Consistent with this possibility, the heterozygous hermaphrodites for the *ego-1(S1198L)*, derived from homozygous hermaphrodites (Fig. EV2A, B), and wild-type (*tmC18[tmIs1200]/+*) males displayed the Rde phenotype. In contrast, heterozygotes for the *ego-1* deletion mutant allele, *ego-1(tm521)*, showed a normal germline exo-RNAi response (Fig. EV2A, B). This dominant effect was also observed in *ego-1(R539Q)*, but not in *ego-1*(V1128E) or *ego-1(C823Y)* heterozygotes (Fig. EV2D, E). We also found that *ego-1(*S1198L*)/ego-1(tm521)* trans-heterozygotes, derived from *ego-1(*S1198L) homozygous hermaphrodites and *ego-1(tm521)* heterozygous males, exhibited a slight reduction in brood size at 25°C, comparable to that of *ego-1(*S1198L) heterozygotes (Fig. EV2C). In addition, they displayed the Rde phenotype when subjected to *pos-1* and *pop-1* RNAi (Fig. EV2D, E). These data suggest that the observed phenotypes of *ego-1(*S1198L) heterozygotes thus far resulted from antimorphic and/or epigenetic effects (see below) rather than haploinsufficiency at the *ego-1* locus.

### Disconnection between *ego-1* genotype and phenotype across generations

Disruption of P granule formation results in aberrant siRNA expression and abnormal suppression of germline RNAi genes, including *sid-1* and *rde-11*, over several generations (Dodson & Kennedy, 2019; Ouyang *et al*., 2019). To determine whether this phenomenon underlies the germline Rde phenotype observed in *ego-1(L1198L)* heterozygotes, we investigated the germline exo-RNAi-defective phenotype over generations following mating with *tmC18[tmIs1200]*, which is a wild-type for the *ego-1* gene. We found that both *ego-1(S1198L)/tmC18[tmIs1200]* heterozygotes and *ego-1(+)* homozygotes (genotype *tmC18[tmIs1200]/tmC18[tmIs1200]*), which originated from *ego-1(S1198L)* heterozygous hermaphrodites derived from *ego-1(S1198L)* homozygous hermaphrodites and wild-type (*tmC18[tmIs1200]*/+) males, exhibited a germline exo-RNAi defective phenotype (Fig. 6A–C). We also found that the germline RNAi-defective phenotype of *ego-1(+)*, originating from *ego-1(S1198L)* heterozygous hermaphrodites, persisted for five generations (Fig. 6B, C), indicating that the germline exo-RNAi defects found in *ego-1(S1198L)/tmC18[tmIs1200]* heterozygotes resulted from epigenetic inheritance initiated by the homozygous *ego-1(S1198L)* mutation. Interestingly, *ego-1(S1198L)/tmC18[tmIs1200]* heterozygotes exhibited more persistent RNAi defects than *ego-1(+)* originating from *ego-1(S1198L)* heterozygous hermaphrodites, suggesting that *ego-1(S1198L)* likely functions predominantly in this context (Fig. 6D–F). We could not determine whether the prolonged persistence of germline exo-RNAi defects in *ego-1(S1198L)* heterozygotes, compared to wild-type animals, was due to the antimorphic or epigenetic effects of the *ego-1(S1198L)* mutation or haploinsufficiency of the *ego-1* gene.

**Figure 6.**
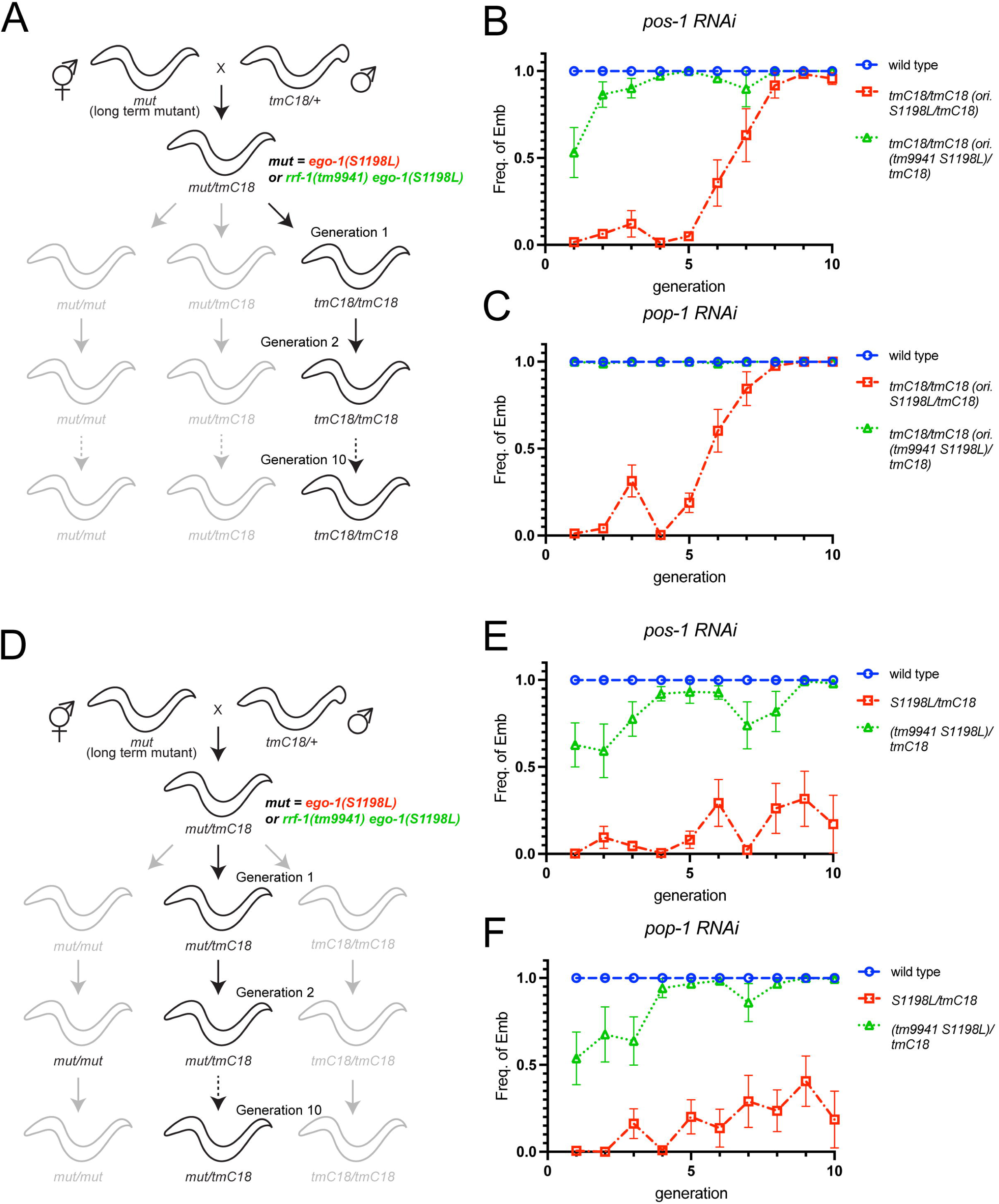
Transgenerational effects of *ego-1(S1198L)* on germline exo-RNAi. A and D. Schematics of genetic crosses. P0 is the first generation in which cross-progenies are selected. B and C. Quantification of the frequency of the indicated mutant animals showing the Emb phenotype under *pos-1* (B) and *pop-1* (C) feeding RNAi in the indicated generation. The wild-type was represented by N2 animals. *tmC18/tmC18* (ori. *S1198L/tmC18*) and *tmC18/tmC18* (ori. *tm9941 S1198L/tmC18*) are descendants of A, where mut represents *ego-1(S1198L)* or *rrf-1(tm9941) ego-1(S1198L)*. E and F. Quantification of the frequency of the indicated mutant animals showing the Emb phenotype under *pos-1* (B) and *pop-1* (C) feeding RNAi in the indicated generation. The wild-type was represented by N2 animals. *S1198L/tmC18* and (*tm9941 S1198L)/tmC18* are the descendants indicated in D, where mut represents *ego-1(S1198L)* or *rrf-1(tm9941) ego-1(S1198L)*.

Given the redundant involvement of EGO-1 and RRF-1 in temperature-dependent sterility, we examined germline exo-RNAi activity in the descendants of *ego-1(S1198L) rrf-1(tm9941)* double-mutant animals. In contrast to *ego-1(+)* animals from *ego-1(S1198L)*, this epigenetic effect was not observed in *rrf-1(+) ego-1(+)* animals from *rrf-1(tm9941) ego-1(S1198L)*, suggesting that ancestral RRF-1 is involved in the establishment of this epigenetic state (Fig. 6B, C). Taken together, these findings highlight the counteractive interplay of RdRPs in the inheritance of germline exo-RNAi-defective phenotypes, along with the redundant enzymatic activity directly required for the silencing of exogenous genes.

### RRF-1 is required for the persistent silencing of *sid-1* and *rde-11* over generations in the descendants of *ego-1(S1198L)*

The *ego-1(S1198L) rrf-1(tm9941)* double mutant itself exhibited germline exo-RNAi defects, but these defects did not persist in the wild-type animals derived from this strain. To investigate how RRF-1 affects the regulation of germline exo-RNAi genes, we analyzed the expression of *sid-1* and *rde-11* in *ego-1(S1198L)* and *rrf-1(tm9951)* single mutants and *ego-1(S1198L) rrf-1(tm9941)* double-homozygous mutants cultured for more than 10 generations. The levels of *sid-1* and *rde-11* expression in *rrf-1(tm9951)* animals were similar to those in wild-type animals (Fig. 7A, B). In *ego-1(S1198L) rrf-1(tm9941)* double-mutant animals, the levels of *sid-1* and *rde-11* were significantly decreased compared to those in the wild-type, but were comparable to those in *ego-1(S1198L)* single-mutant animals (Fig. 7A, B). Next, we examined the expression of *sid-1* and *rde-11* in wild-type descendants derived from *ego-1(S1198L)* and *ego-1(S1198L) rrf-1(tm9941)* mutant animals at generations 3, 5, 7, and 10 after crossing with the *tmC18* balancer (Fig. 3A). In the descendants, the expression level of *sid-1* recovered progressively with each generation when derived from *ego-1(S1198L)* mutants, whereas it recovered in the *ego-1(S1198L) rrf-1(tm9941)* mutant in the early generations (Fig. 7C). The expression level of *rde-11* in *ego-1(S1198L) rrf-1(tm9941)* mutant-derived descendants was higher than that in *ego-1(S1198L)* mutant-derived descendants (Fig. 7D). These results suggest that EGO-1(S1198L) is defective in licensing the expression of *sid-1* and *rde-11*. In contrast, RRF-1 mediated the silencing of *sid-1* and *rde-11* when EGO-1 function was inhibited.

**Figure 7.**
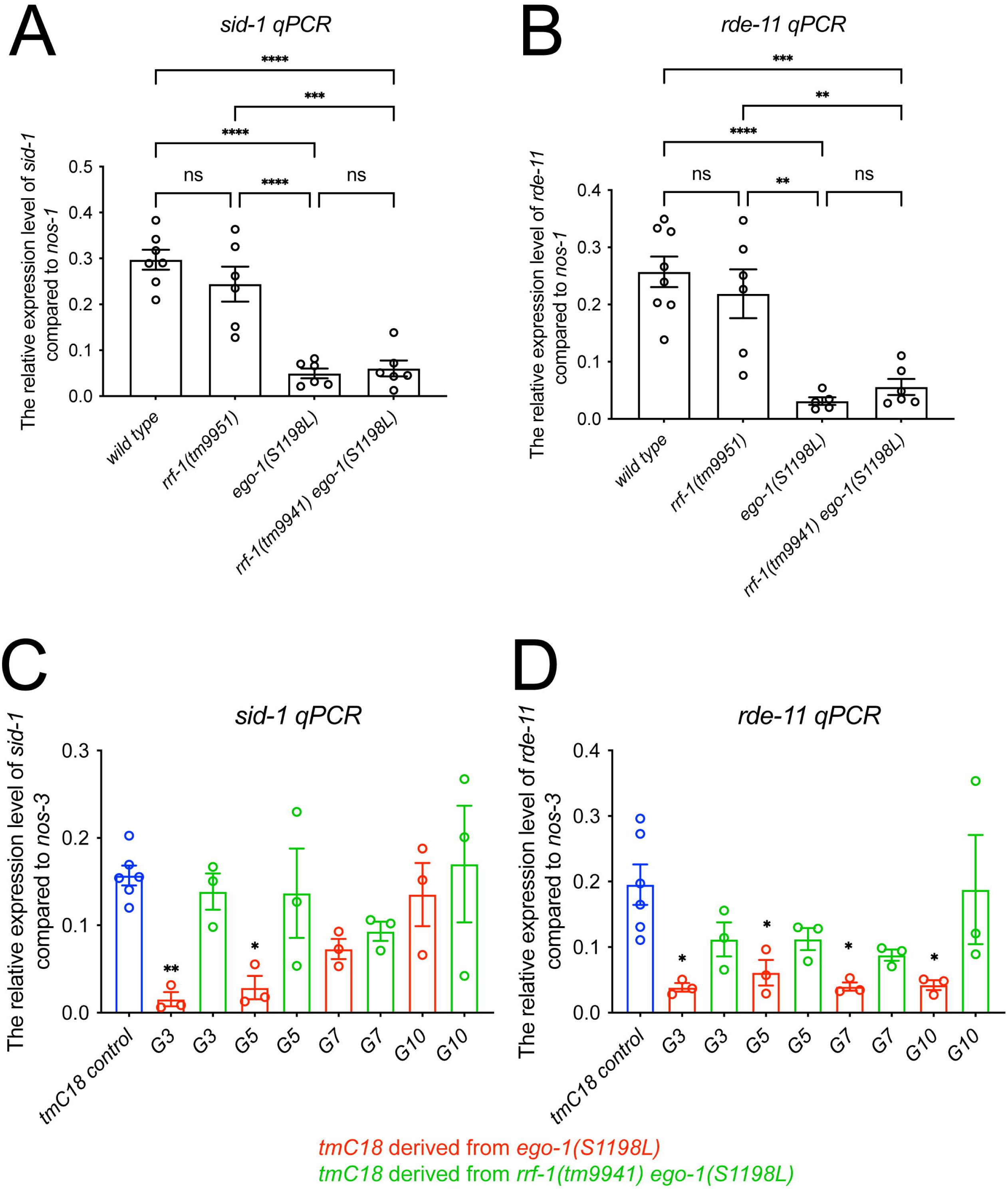
RRF-1-dependent regulation of *sid-1* and *rde-11* expression across generations in *ego-1(S1198L)* mutants. A and B. RT-qPCR showing the relative expression levels of *sid-1* (A) and *rde-11* (B) mRNA normalized to *nos-3* expression. RNA was extracted from young-adult animals grown at 20°C. C and D. RT-qPCR showing the relative expression levels of *sid-1* (C) and *rde-11* (D) mRNA normalized to *nos-3* expression. RNA was extracted from young-adult animals grown at 20°C.

## Discussion

Investigating the unique roles of EGO-1 and RRF-1 is valuable, as it helps explain how *C. elegans* manages gene expression through different RNAi mechanisms, including those initiated by exogenous dsRNA. Although it has been suggested that these molecules may have overlapping functions in exo-RNAi(Sijen *et al*., 2001; Smardon *et al*., 2000), the underlying mechanisms remain unclear. Previous studies have reported that P granules protect *sid-1* and *rde-11* mRNAs from PRG-1/HRDE-1-dependent silencing(Dodson & Kennedy, 2019; Lev *et al*., 2019; Ouyang *et al*., 2019). Additionally, CSR-1 protects its targets from silencing by PRG-1(Seth *et al*, 2013; Wedeles *et al*, 2013). Given that EGO-1 and RRF-1 are responsible for the synthesis of CSR-1-class and WAGO-class 22G RNAs, respectively, a model can be considered in which RRF-1 amplifies 22G RNAs in PRG-1/HRDE-1-dependent silencing, whereas EGO-1 protects targets from PRG-1/HRDE-1-mediated silencing (Fig. 8). However, testing this hypothesis is challenging because *ego-1* null mutants are sterile. In this study, we provide experimental evidence supporting this hypothesis. We analyzed *ego-1* alleles in the MMP collection, which revealed germline exo-RNAi defects in *ego-1(S1198L)*. Despite exhibiting these defects, this allele retains its fundamental function in temperature-independent germline development, providing valuable insights into the specific role of EGO-1 in RNAi mechanisms (Fig. 8). Additionally, this allele showed synthetic effects with the *rrf-1* deletion mutation on temperature-dependent sterility. Germline exo-RNAi defects in *ego-1(S1198L)* were partially suppressed by the inhibition of HRDE-1, CDE-1, and ZNFX-1, whereas inheritance of the RNAi-defective phenotype across generations occurred in an RRF-1-dependent manner. Thus, while EGO-1 functions redundantly with RRF-1 to undergo temperature-dependent gametogenesis, EGO-1 and RRF-1 counteract each other during HRDE-1-dependent germline gene silencing. Our study highlights the dual role of EGO-1 in enhancing germline robustness to exo-RNAi via RdRP activity, which is directly required for target gene silencing and protecting exo-RNAi genes from HRDE-1-dependent silencing.

**Figure 8.**
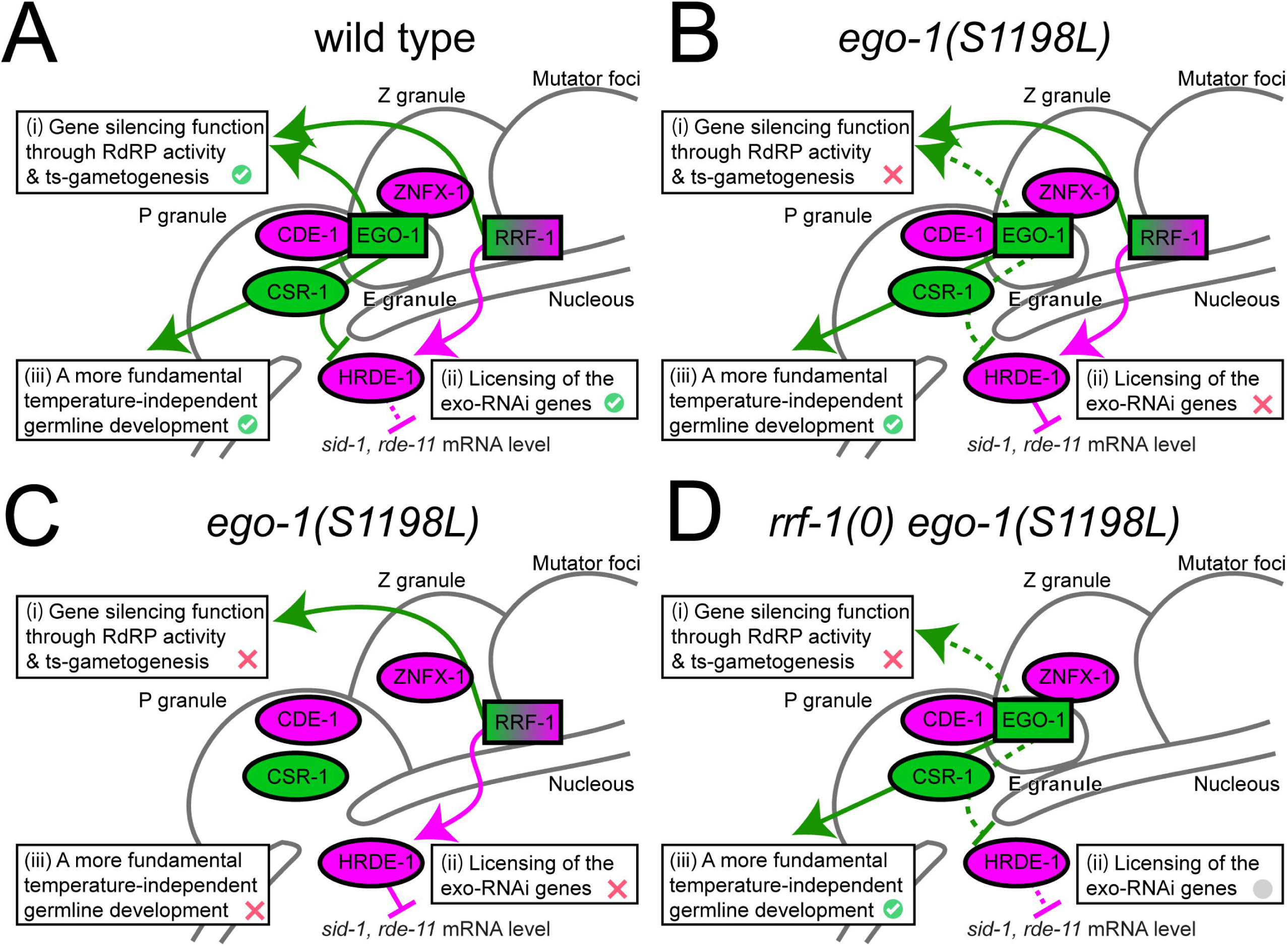
Model for the regulation of germline exo-RNAi by EGO-1 and RRF-1. A. As noted in previous studies, EGO-1 and RRF-1 share a common role in gene silencing via RdRP activity in germline exoRNAi (i). However, these RdRPs play distinct roles in the epigenetic regulation of exo-RNAi gene expression (ii). EGO-1 represses the transcriptional inhibition of exo-RNAi genes by HRDE-1, probably through the synthesis of CSR-1 class 22G RNA. In contrast, RRF-1 is responsible for the synthesis of WAGO class 22G RNAs loaded with HRDE-1 and positively acts against the transcriptional inhibition of exo-RNAi genes. Additionally, EGO-1, but not RRF-1, plays an essential role in fundamental temperature-independent germline development, including heterochromatin assembly (iii) (Maine *et al*, 2005). B. The *ego-1(S1198L)* mutation reduced the inhibitory activity against HRDE-1 (green dotted T-line), and RdRP activity was directly required for exo-RNAi-mediated gene silencing (arrow with green dotted line). Some functions specific to EGO-1, including its fundamental role in temperature-independent germline development, remained unchanged in this mutant. C. In the *ego-1* null mutants, both the inhibitory activity against HRDE-1 and the RdRP activity directly required for exo-RNAi-mediated gene silencing were lost. D. In the double mutant germline of *rrf-1(0)* and *ego-1(S1198L)*, HRED-1-dependent repression of germline exo-RNAi genes did not function well. However, RdRP, which is directly required for exo-RNAi-mediated gene silencing, was inactivated, resulting in germline exo-RNAi defects.

It is known that *ego-1* is involved in both germline exo-RNAi and germline development; however, it is not clear whether germline abnormalities and exo-RNAi defects occur independently or concurrently. Here, we found that some *ego-1* mutants were fertile, despite germline exo-RNAi defects. In the *ego-1(S1198L)* mutant, germline exo-RNAi defects were observed without reproductive developmental abnormalities, but abnormal distributions of PGL-1 and ZNFX-1 were observed. This suggests that the inhibition of EGO-1 does not necessarily cause RNAi defects due to reproductive developmental abnormalities; rather, the abnormal function of P granules and/or Z granules may contribute to RNAi defects, along with the RdRP activity of EGO-1, which is directly required for gene silencing.

Mutations with germline exo-RNAi defects have amino acid substitutions in the residues that are predicted as “deleterious” mutations by the PROVEAN and PolyPhen-2 prediction programs. C823Y and R539Q are substitutions located within the RdRP domain but in close proximity, whereas V1128E and S1198L are substitutions located outside the RdRP domain. S1198 and R539, whose mutations are associated with strong germline exo-RNAi defects, were predicted to be inside the protein using Alpha Fold 3D structure prediction. These mutations are relatively close to each other: the distance between R539 and S1198 is 15.1 Å, between C823 and S1198 is 32.49 Å, and between V1128 and S1198 is 29.64 Å. This proximity raises the possibility that these mutations may similarly affect RdRP activity and/or influence interactions with other molecules, such as CDE-1 and ZNFX-1, through allosteric effects. It is intriguing that these four mutations correspond to amino acids conserved in RRF-1. Future studies, including chimeric experiments involving EGO-1 and RRF-1, will provide more detailed insights into the underlying mechanisms. The robustness of the network regulating RNAi is crucial not only for biological processes, but also as an experimental tool for gene knockdown. Further elucidation of the working mechanisms of RdRPs could shed light on the developmental and antiviral roles of RNAi and broaden its technological applications.

## Structured Methods

### Reagents and Tools Table

**Structured Methods - Reagents and Tools Table**
| Reagent/Resource | Reference or Source | Identifier or Catalog Number |
| --- | --- | --- |
| **Experimental Models** |  |  |
| *ego-1(gk540555) I* | VC40259 was outcrossed with *tmC18[tmls1200]* once | FX31713 |
| *ego-1(gk864727) I* | VC40886 was outcrossed with *tmC18[tmls1200]* once | FX31714 |
| *ego-1(gk115390) I* | VC20206 was outcrossed with *tmC18[tmls1200]* once | FX31715 |
| *ego-1(gk317493) I* | VC20319 was outcrossed with *tmC18[tmls1200]* once | FX31716 |
| *ego-1(gk115391) I* | VC30058 was outcrossed with *tmC18[tmls1200]* once | FX31717 |
| *ego-1(gk532049) I* | VC40244 was outcrossed with *tmC18[tmls1200]* once | FX31718 |
| *ego-1(gk749674) I* | VC40660 was outcrossed with *tmC18[tmls1200]* once | FX31719 |
| *ego-1(gk925207) I* | VC41006 was outcrossed with *tmC18[tmls1200]* once | FX31720 |
| *ego-1(gk357146) I* | VC20618 was outcrossed with *tmC18[tmls1200]* once | FX31721 |
| *ego-1(gk721963) I* | VC40613 was outcrossed with *tmC18[tmls1200]* once | FX31722 |
| *ego-1(gk882383) I* | VC40920 was outcrossed with *tmC18[tmls1200]* once | FX31723 |
| *ego-1(gk481348) I* | VC40140 was outcrossed with *tmC18[tmls1200]* once | FX31724 |
| *ego-1(gk115393) I* | VC20439 was outcrossed with *tmC18[tmls1200]* once | FX31725 |
| *ego-1(gk896494) I* | VC40951 was outcrossed with *tmC18[tmls1200]* once | FX31726 |
| *ego-1(gk426642) I* | VC30158 was outcrossed with *tmC18[tmls1200]* once | FX31727 |
| *ego-1(gk115394) I* | VC20474 was outcrossed with *tmC18[tmls1200]* once | FX31728 |
| *ego-1(gk837385) I* | VC40832 was outcrossed with *tmC18[tmls1200]* once | FX31729 |
| *ego-1(gk115395) I* | VC40084 was outcrossed with *tmC18[tmls1200]* once | FX31730 |
| *ego-1(gk115397) I* | VC20545 was outcrossed with *tmC18[tmls1200]* once | FX31731 |
| *ego-1(gk498425) I* | VC40175 was outcrossed with *tmC18[tmls1200]* once | FX31732 |
| *ego-1(gk470185) I* | VC40116 was outcrossed with *tmC18[tmls1200]* once | FX31733 |
| *ego-1(gk720210) I* | VC40611 was outcrossed with *tmC18[tmls1200]* once | FX31734 |
| *ego-1(gk115401) I* | VC40050 was outcrossed with *tmC18[tmls1200]* once | FX31735 |
| *ego-1(gk532049) I* | This study. VC40244 was outcrossed with *tmC18[tmls1200]* 4 times | FX34341 |
| *ego-1(gk426642) I* | This study. VC30158 was outcrossed with *tmC18[tmls1200]* 4 times | FX34380 |
| *ego-1(gk882383) I* | This study. VC40920 was outcrossed with *tmC18[tmls1200]* 4 times | FX34343 |
| *ego-1(gk357146) I* | This study. VC20618 was outcrossed with *tmC18[tmls1200]* 4 times | FX34342 |
| *ego-1(gk115390) I* | This study. VC20206 was outcrossed with *tmC18[tmls1200]* 4 times | FX34340 |
| *rrf-1(pk1417) I* | from CGC | MAH23 |
| *rrf-1(tm9951) I* | This study, created by CRISPR-Cas9. F26A3: 6169/6170-7212/7213(1043 bp deletion) | FX9951 |
| *rrf-1(tm9941)*<br>*ego-1(gk532049) I* | This study, created by CRISPR-Cas9. F26A3:5756/5757-7220/7221 (1464 bp deletion) | FX9941 |
| *mut-15(tm1358) V* | from NBRP | FX01358 |
| *mut-14(pk738) V* | from CGC | NL1838 |
| *tmC18[tmls1200] I* | from NBRP | FX30167 |
| *ego-1(tm521)/tmC18[tmls1200] I* | from NBRP | FX33930 |
| *cde-1(tm1021) III* | FX01021 *cde-1(tm1021)* was outcrossed with N2 3 times, from NBRP | FX34683 |
| *ego-1(gk532049) I*;<br>*cde-1(tm1021) III* | This study | FX34682 |
| *hrde-1(tm1200) III* | FX01200 *hrde-1(tm1200)* was outcrossed with N2 twice, from NBRP | FX34449 |
| *ego-1(gk532049) I*;<br>*hrde-1(tm1200) III* | This study | FX34348 |
| *hrde-1(tor125[hrde-1::gfp]) III* | from CGC | JMC231 |
| *ego-1(gk532049)/tmC18[tmls1200] I*; *hrde-1::gfp(tor125) III* | This study | FX34770 |
| *znfx-1(gg544[3xflag::gfp::znfx-1]) II*; | from CGC | YY968 |
| *pgl-1(gg547[pgl-1::3xflag::tag RFP]) IV* |  |  |
| *ego-1(gk532049)/tmC18[tmls1 236] I;*<br>*znfx-1(gg544[3xflag::gfp::znfx-1]) II;*<br>*pgl-1(gg547[pgl-1::3xflag::tag RFP]) IV* | This study | FX34771 |
| *znfx-1(gg561) II* | from CGC | YY996 |
| *ego-1(gk532049)/tmC18[tmls1 200] I; znfx-1(gg561) II* | This study | FX34772 |
| *C. elegans* wild type | CGC | N2 |
| **Recombinant DNA** |  |  |
| rrf-1_sg1_pDD162 | This study | rrf-1_sg1 |
| rrf-2_sg1_pDD162 | This study | rrf-1_sg2 |
| pDD162 | Addgene | Plasmid #47549 |
| pCFJ90 | Addgene | Plasmid #19327 |
| **Oligonucleotides and other sequence-based reagents** |  |  |
| primers used for construction of CRISPR plasmids | Eurofins | Methods and Protocols |
| primers used for quantitative PCR | Eurofins | Methods and Protocols |
| **Chemicals, Enzymes and other reagents** |  |  |
| isopropyl β-D-1-thiogalactopyranoside (IPTG) | Sigma Aldrich | I6758 |
| TRIzol | Invitrogen | Cat: 15596026 |
| SuperScript IV Reverse Transcriptase | Invitrogen | Cat: 18090010 |
| Power™ SYBR™ Green | Applied Biosystems | Cat: 4368702 |
| Master Mix |  |  |
| **Software** |  |  |
| ImageJ | (Schneider *et al*, 2012) | SCR_003070 |
| Prism GraphPad 10 for MacOS X | GrapPad Software | N/A |

### Methods and Protocols

#### Nematode culture and strains

Unless otherwise noted, worms were grown under standard conditions at 20°C. The wild-type strain was Bristol N2. The food source used was *Escherichia coli* strain OP50-1. The *C. elegans* strain information is summarized in Reagents and Tools Table.

#### CRISPR/Cas9-mediated genome editing

To generate deletion mutations in the *rrf-1* gene, we used the plasmid-based CRISPR/Cas9 method as described previously (Dejima *et al*, 2023; Dickinson *et al*, 2013). We injected a genome editing plasmid mixture containing two Cas9/sgRNA plasmids targeting *rrf-1* (25 ng/μl each), an injection marker plasmid pCFJ90 (Pmyo-2::mCherry) (5 ng/μl), and an NEB 1 kb DNA ladder (145 ng/μl) into the gonads of wild-type or *ego-1(gk532049)* animals. The deletions were identified by PCR screening and confirmed by Sanger sequencing. The primers used in Cas9/sgRNA plasmid construction are:

rrf-1_sg1Fwd 5’-TACATCGTCTCCGTCTCGTTTTAGAGCTAGAAATAGCAAGT-3’
rrf-1_sg1Rev 5’-GACGGAGACGATGTAACGCAAGACATCTCGCAATAGG-3’
rrf-1_sg2Fwd 5’-AACACAGAGTCCAACCCGTTTTAGAGCTAGAAATAGCAAGT-3’
rrf-1_sg2Rev 5’-GTTGGACTCTGTGTTCTTCAAGACATCTCGCAATAGG-3’.

#### Outcrossing with a balancer

The *ego-1* mutants were obtained from the Caenorhabditis elegans Genetics Center, which is supported by the National Institutes of Health National Center for Research Resources. To outcross the strains, males heterozygous for *tmC18[tmIs1200]* were crossed with each *ego-1* mutant strain. GFP-positive *tmC18[tmIs1200]* was singled over mutant trans-heterozygous F1 hermaphrodites, and their GFP-negative mutant homozygous progeny (F2) was further singled and propagated.

#### Quantitative PCR

Total RNA was isolated from 20–30 day-1 adults per replicate using the TRIzol reagent (Invitrogen). Superscript IV reverse transcriptase (Invitrogen) was used for reverse transcription according to the manufacturer’s instructions. DNase I-treated total RNA (100 ng) was used for the reverse transcription reaction, and 1/10 of the reverse transcription product was used as the template for the PCR reaction. Quantitative PCR was performed in a 7500 Real-time Thermal cycler (Applied Biosystems) using the Power SYBR master mix (Applied Biosystems) with the following parameters: 95°C for 10 min, and 40 cycles of 95°C for 15 s and 55°C for 60 s. Data were normalized to the *nos-3* gene. Gonads were atrophied in the *ego-1* null mutant. Therefore, *ama-1* was used for normalization instead of *nos-3*.

The primers used are the same primers as those used in (Dejima *et al*., 2023; Dodson & Kennedy, 2019):

*nos-3*
Y53C12B#F17 5’-GGAGGCTATCGGCAGTATCA-3’
Y53C12B#R19 5’-GTGGCCCTGCTTGAGGATTA-3’
*rde-11*
B0564#F22 5’-GATTTCGGACTCCCTATGTGGAC-3’
B0564#R21 5’-GTAGAGATACAGTCCGTCCAGC-3’
*sid-1*
C04F5#F35 5’-CGGCGAATGAATCCATCTAT-3’
C04F5#R20 5’-CGGGAGCTATGAAGACGAAG-3’
*ama-1*
ama-1_Q#F2 5’-AGATGGACCTCACCGACAAC-3’
ama-1_Q#R2 5’-CTGCAGATTACACGGAAGCA-3’

#### RNAi by feeding method

RNAi was performed as described previously (Dejima *et al*., 2023; Yoshida *et al*, 2023). For *pos-1* and *pop-1* feeding RNAi, the L4 worms were cultured on RNAi plates for 24 h. Several adult animals were transferred to new NGM plates, allowed to lay eggs for several hours, and then removed. The percentage of dead eggs was calculated after 24 h.

#### Image analysis and quantification

Animals expressing fluorescent proteins were mounted on 5% agar pads in the presence of 10–25 mM NaN_3_ and imaged with a BX51 microscope equipped with a DP80 CCD camera (Olympus Optical Co., Ltd.). Images were captured at 40× magnification with a resolution of 1360×1024 pixels, and identical settings and exposure times were applied for each fluorescent protein. Fluorescence intensity quantification was performed using custom image analysis scripts implemented in Fiji. To determine the regions of interest (ROIs) for the particles of GFP::ZNFX-1 and tagRFP::PGL-1, the following processing steps were performed. A duplicate was created for each image, the Fast Fourier Transform (FFT) was applied, and a rectangular region in the center of the FFT-applied image with dimensions of 30×30 pixels was cleared. The resulting image, resembling noise-reduced images perceptible to the human eye, was obtained by applying inverse FFT to transform the image back into the spatial domain. Then, background subtraction (rolling ball algorithm with a rolling radius of 50 pixels), thresholding (Huang dark method), and morphological operations (erosion and watershed) were employed to segment individual puncta. The resulting ROIs were then applied to the original image for intensity measurement. To analyze HRDE-1::GFP, a single nucleus with the optimal focus within the pachytene region was arbitrarily chosen by selecting a square region (104×104 pixels) per animal. A rolling ball algorithm with a rolling radius of 20 pixels was applied to subtract background fluorescence. Then, automatic thresholding using the Huang method was performed to convert the image into a binary representation, highlighting the ROI. The image was converted to a binary mask to isolate the nucleus from the background. A selection corresponding to the segmented nucleus was created for subsequent intensity measurements. This selection was then applied to the original image, and the fluorescence intensity in the nucleus was quantified. Cytosolic intensity was calculated by subtracting the nuclear intensity from the overall intensity.

### Statistics

Statistical analyses were performed using Prism 10 (GraphPad Software, Inc.). All bar graphs show the means with error bars (with ± SEM). Comparisons between more than two groups were performed using one-way ANOVA. Tukey’s multiple-comparisons test was used for multiple comparisons. Data were considered statistically significant at a p-value of less than 0.05. * indicates pL<L0.05, ** indicates pL<L0.01, *** indicates p < 0.005, and **** indicates <0.001.

## Acknowledgments

We thank the members of the Mitani Laboratory for their support. Some strains were provided by the CGC, which was funded by the NIH Office of Research Infrastructure Programs (P40 OD010440). This work was supported by Technology of Japan Grants-in-Aid for Scientific Research to KD (20K06561, 24K09370) and SM (20H03422), and a grant from the Takeda Science Foundation.

## Author contributions

Katsufumi Dejima: Conceptualization; Funding acquisition; Investigation; Methodology; Writing – original draft; Writing – review and editing. Keita Yoshida: Investigation; Writing – review and editing. Shohei Mitani: Conceptualization; Supervision; Funding acquisition; Writing – original draft; Project administration; Writing – review and editing.

## Data availability

All data generated or analyzed in this study are included in the published article and its supplementary information files.

**Figure EV1.** *ego-1(S1198L)* is sensitive to somatic RNAi. A and B. Quantification of the frequency of the indicated mutant animals showing the expected phenotype under *bli-3* (A) and *bli-1* (B) feeding RNAi.

**Figure EV2.** RNAi-defective phenotype caused by *ego-1(R539Q)* and *ego-1(S1198L)* heterozygous animals. A. Quantification of the frequency of indicated mutant animals showing the Emb phenotype under *pos-1* feeding RNAi. B. Quantification of the frequency of indicated mutant animals showing the Emb phenotype under *pop-1* feeding RNAi. C. Brood size analysis of the indicated mutant animals that were cultured at 25°C. D. Quantification of the frequency of indicated mutant animals showing the Emb phenotype under *pos-1* feeding RNAi. E. Quantification of the frequency of indicated mutant animals showing the Emb phenotype under *pop-1* feeding RNAi.

## References

Adzhubei IA, Schmidt S, Peshkin L, Ramensky VE, Gerasimova A, Bork P, Kondrashov AS, Sunyaev SR (2010) A method and server for predicting damaging missense mutations. Nat Methods 7: 248–249

Ashe A, Sapetschnig A, Weick EM, Mitchell J, Bagijn MP, Cording AC, Doebley AL, Goldstein LD, Lehrbach NJ, Le Pen J et al (2012) piRNAs can trigger a multigenerational epigenetic memory in the germline of C. elegans. Cell 150: 88–99

Batista PJ, Ruby JG, Claycomb JM, Chiang R, Fahlgren N, Kasschau KD, Chaves DA, Gu W, Vasale JJ, Duan S et al (2008) PRG-1 and 21U-RNAs interact to form the piRNA complex required for fertility in C. elegans. Molecular cell 31: 67–78

Buckley BA, Burkhart KB, Gu SG, Spracklin G, Kershner A, Fritz H, Kimble J, Fire A, Kennedy S (2012) A nuclear Argonaute promotes multigenerational epigenetic inheritance and germline immortality. Nature 489: 447–451

Castel SE, Martienssen RA (2013) RNA interference in the nucleus: roles for small RNAs in transcription, epigenetics and beyond. Nat Rev Genet 14: 100–112

Cecere G, Zheng GX, Mansisidor AR, Klymko KE, Grishok A (2012) Promoters recognized by forkhead proteins exist for individual 21U-RNAs. Molecular cell 47: 734–745

Chen X, Wang K, Mufti FUD, Xu D, Zhu C, Huang X, Zeng C, Jin Q, Huang X, Yan YH et al (2024) Germ granule compartments coordinate specialized small RNA production. Nat Commun 15: 5799

Choi Y, Sims GE, Murphy S, Miller JR, Chan AP (2012) Predicting the functional effect of amino acid substitutions and indels. PLoS One 7: e46688

Claycomb JM, Batista PJ, Pang KM, Gu W, Vasale JJ, van Wolfswinkel JC, Chaves DA, Shirayama M, Mitani S, Ketting RF et al (2009) The Argonaute CSR-1 and its 22G-RNA cofactors are required for holocentric chromosome segregation. Cell 139: 123–134

Conine CC, Batista PJ, Gu W, Claycomb JM, Chaves DA, Shirayama M, Mello CC (2010) Argonautes ALG-3 and ALG-4 are required for spermatogenesis-specific 26G-RNAs and thermotolerant sperm in Caenorhabditis elegans. Proc Natl Acad Sci U S A 107: 3588–3593

Das PP, Bagijn MP, Goldstein LD, Woolford JR, Lehrbach NJ, Sapetschnig A, Buhecha HR, Gilchrist MJ, Howe KL, Stark R et al (2008) Piwi and piRNAs act upstream of an endogenous siRNA pathway to suppress Tc3 transposon mobility in the Caenorhabditis elegans germline. Molecular cell 31: 79–90

Dejima K, Imae R, Suehiro Y, Yoshida K, Mitani S (2023) An endomembrane zinc transporter negatively regulates systemic RNAi in Caenorhabditis elegans. iScience 26: 106930

Dejima K, Mitani S (2022) Balancer-assisted outcrossing to remove unwanted background mutations. MicroPubl Biol 2022

Dickinson DJ, Ward JD, Reiner DJ, Goldstein B (2013) Engineering the *Caenorhabditis elegans* genome using Cas9-triggered homologous recombination. Nat Methods 10: 1028–1034

Dodson AE, Kennedy S (2019) Germ Granules Coordinate RNA-Based Epigenetic Inheritance Pathways. Dev Cell 50: 704–715 e704

Fire A, Xu S, Montgomery MK, Kostas SA, Driver SE, Mello CC (1998) Potent and specific genetic interference by double-stranded RNA in *Caenorhabditis elegans*. Nature 391: 806–811

Gent JI, Lamm AT, Pavelec DM, Maniar JM, Parameswaran P, Tao L, Kennedy S, Fire AZ (2010) Distinct phases of siRNA synthesis in an endogenous RNAi pathway in C. elegans soma. Molecular cell 37: 679–689

Gu W, Shirayama M, Conte D, Jr., Vasale J, Batista PJ, Claycomb JM, Moresco JJ, Youngman EM, Keys J, Stoltz MJ et al (2009) Distinct argonaute-mediated 22G-RNA pathways direct genome surveillance in the C. elegans germline. Molecular cell 36: 231–244

Ishidate T, Ozturk AR, Durning DJ, Sharma R, Shen EZ, Chen H, Seth M, Shirayama M, Mello CC (2018) ZNFX-1 Functions within Perinuclear Nuage to Balance Epigenetic Signals. Molecular cell 70: 639–649 e636

Kennedy S, Wang D, Ruvkun G (2004) A conserved siRNA-degrading RNase negatively regulates RNA interference in *C. elegans*. Nature 427: 645–649

Ketting RF, Haverkamp TH, van Luenen HG, Plasterk RH (1999) Mut-7 of C. elegans, required for transposon silencing and RNA interference, is a homolog of Werner syndrome helicase and RNaseD. Cell 99: 133–141

Kumsta C, Hansen M (2012) C. elegans rrf-1 mutations maintain RNAi efficiency in the soma in addition to the germline. PLoS One 7: e35428

Lev I, Toker IA, Mor Y, Nitzan A, Weintraub G, Antonova O, Bhonkar O, Ben Shushan I, Seroussi U, Claycomb JM et al (2019) Germ Granules Govern Small RNA Inheritance. Curr Biol 29: 2880–2891 e2884

Luteijn MJ, van Bergeijk P, Kaaij LJ, Almeida MV, Roovers EF, Berezikov E, Ketting RF (2012) Extremely stable Piwi-induced gene silencing in Caenorhabditis elegans. EMBO J 31: 3422–3430

Maine EM, Hauth J, Ratliff T, Vought VE, She X, Kelly WG (2005) EGO-1, a putative RNA-dependent RNA polymerase, is required for heterochromatin assembly on unpaired dna during C. elegans meiosis. Curr Biol 15: 1972–1978

Ng PC, Henikoff S (2001) Predicting deleterious amino acid substitutions. Genome Res 11: 863–874

Ouyang JPT, Folkmann A, Bernard L, Lee CY, Seroussi U, Charlesworth AG, Claycomb JM, Seydoux G (2019) P Granules Protect RNA Interference Genes from Silencing by piRNAs. Dev Cell 50: 716–728 e716

Ouyang JPT, Zhang WL, Seydoux G (2022) The conserved helicase ZNFX-1 memorializes silenced RNAs in perinuclear condensates. Nat Cell Biol 24: 1129–1140

Pavelec DM, Lachowiec J, Duchaine TF, Smith HE, Kennedy S (2009) Requirement for the ERI/DICER complex in endogenous RNA interference and sperm development in Caenorhabditis elegans. Genetics 183: 1283–1295

Phillips CM, Montgomery TA, Breen PC, Ruvkun G (2012) MUT-16 promotes formation of perinuclear mutator foci required for RNA silencing in the *C. elegans* germline. Genes & development 26: 1433–1444

Phillips CM, Updike DL (2022) Germ granules and gene regulation in the Caenorhabditis elegans germline. Genetics 220

Schneider CA, Rasband WS, Eliceiri KW (2012) NIH Image to ImageJ: 25 years of image analysis. Nat Methods 9: 671–675

Seth M, Shirayama M, Gu W, Ishidate T, Conte D, Jr., Mello CC (2013) The C. elegans CSR-1 argonaute pathway counteracts epigenetic silencing to promote germline gene expression. Dev Cell 27: 656–663

Shen EZ, Chen H, Ozturk AR, Tu S, Shirayama M, Tang W, Ding YH, Dai SY, Weng Z, Mello CC (2018) Identification of piRNA Binding Sites Reveals the Argonaute Regulatory Landscape of the C. elegans Germline. Cell 172: 937–951 e918

Shirayama M, Seth M, Lee HC, Gu W, Ishidate T, Conte D, Jr., Mello CC (2012) piRNAs initiate an epigenetic memory of nonself RNA in the C. elegans germline. Cell 150: 65–77

Sijen T, Fleenor J, Simmer F, Thijssen KL, Parrish S, Timmons L, Plasterk RH, Fire A (2001) On the role of RNA amplification in dsRNA-triggered gene silencing. Cell 107: 465–476

Simmer F, Tijsterman M, Parrish S, Koushika SP, Nonet ML, Fire A, Ahringer J, Plasterk RH (2002) Loss of the putative RNA-directed RNA polymerase RRF-3 makes *C. elegans* hypersensitive to RNAi. Curr Biol 12: 1317–1319

Smardon A, Spoerke JM, Stacey SC, Klein ME, Mackin N, Maine EM (2000) EGO-1 is related to RNA-directed RNA polymerase and functions in germ-line development and RNA interference in *C. elegans*. Curr Biol 10: 169–178

Tabara H, Sarkissian M, Kelly WG, Fleenor J, Grishok A, Timmons L, Fire A, Mello CC (1999) The rde-1 gene, RNA interference, and transposon silencing in C. elegans. Cell 99: 123–132

Tabara H, Yigit E, Siomi H, Mello CC (2002) The dsRNA binding protein RDE-4 interacts with RDE-1, DCR-1, and a DExH-box helicase to direct RNAi in C. elegans. Cell 109: 861–871

Thompson O, Edgley M, Strasbourger P, Flibotte S, Ewing B, Adair R, Au V, Chaudhry I, Fernando L, Hutter H et al (2013) The million mutation project: a new approach to genetics in *Caenorhabditis elegans*. Genome Res 23: 1749–1762

van Wolfswinkel JC, Claycomb JM, Batista PJ, Mello CC, Berezikov E, Ketting RF (2009) CDE-1 affects chromosome segregation through uridylation of CSR-1-bound siRNAs. Cell 139: 135–148

Vought VE, Ohmachi M, Lee MH, Maine EM (2005) EGO-1, a putative RNA-directed RNA polymerase, promotes germline proliferation in parallel with GLP-1/notch signaling and regulates the spatial organization of nuclear pore complexes and germline P granules in Caenorhabditis elegans. Genetics 170: 1121–1132

Wang G, Reinke V (2008) A C. elegans Piwi, PRG-1, regulates 21U-RNAs during spermatogenesis. Curr Biol 18: 861–867

Wedeles CJ, Wu MZ, Claycomb JM (2013) Protection of germline gene expression by the C. elegans Argonaute CSR-1. Dev Cell 27: 664–671

Xu F, Feng X, Chen X, Weng C, Yan Q, Xu T, Hong M, Guang S (2018) A Cytoplasmic Argonaute Protein Promotes the Inheritance of RNAi. Cell Rep 23: 2482–2494

Yigit E, Batista PJ, Bei YX, Pang KM, Chen CCG, Tolia NH, Joshua-Tor L, Mitani S, Simard MJ, Mello CC (2006) Analysis of the argonaute family reveals that distinct argonautes act sequentially during RNAi. Cell 127: 747–757

Yoshida K, Suehiro Y, Dejima K, Yoshina S, Mitani S (2023) Distinct pathways for export of silencing RNA in Caenorhabditis elegans systemic RNAi. iScience 26: 108067

Zhou X, Feng X, Mao H, Li M, Xu F, Hu K, Guang S (2017) RdRP-synthesized antisense ribosomal siRNAs silence pre-rRNA via the nuclear RNAi pathway. Nature structural & molecular biology 24: 258–269

