## Supplementary figures and images for "Dual roles of EGO-1 and RRF-1 in regulating germline exo-RNAi efficiency in *Caenorhabditis elegans*"

### Fig. EV1

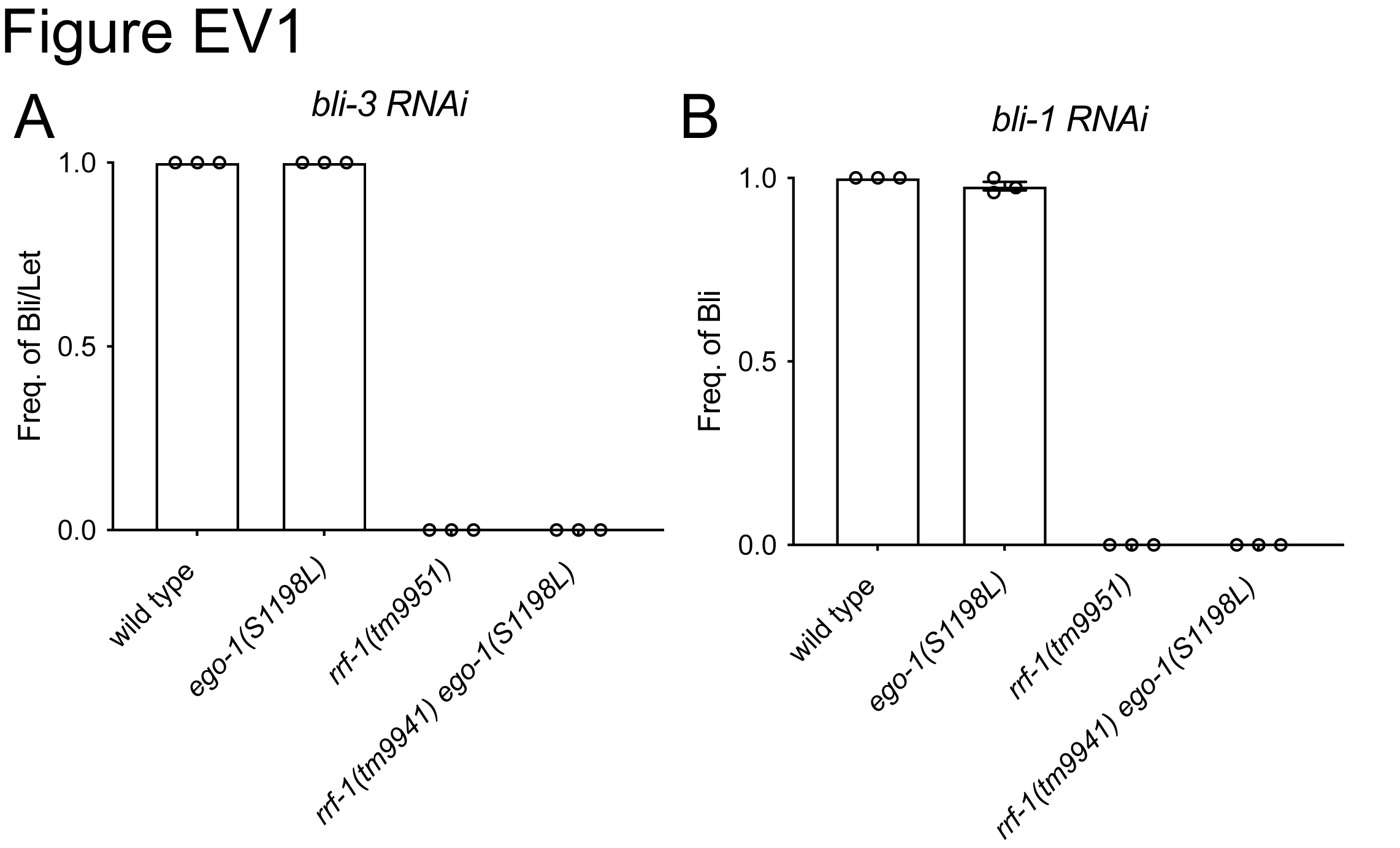

### Fig. EV2

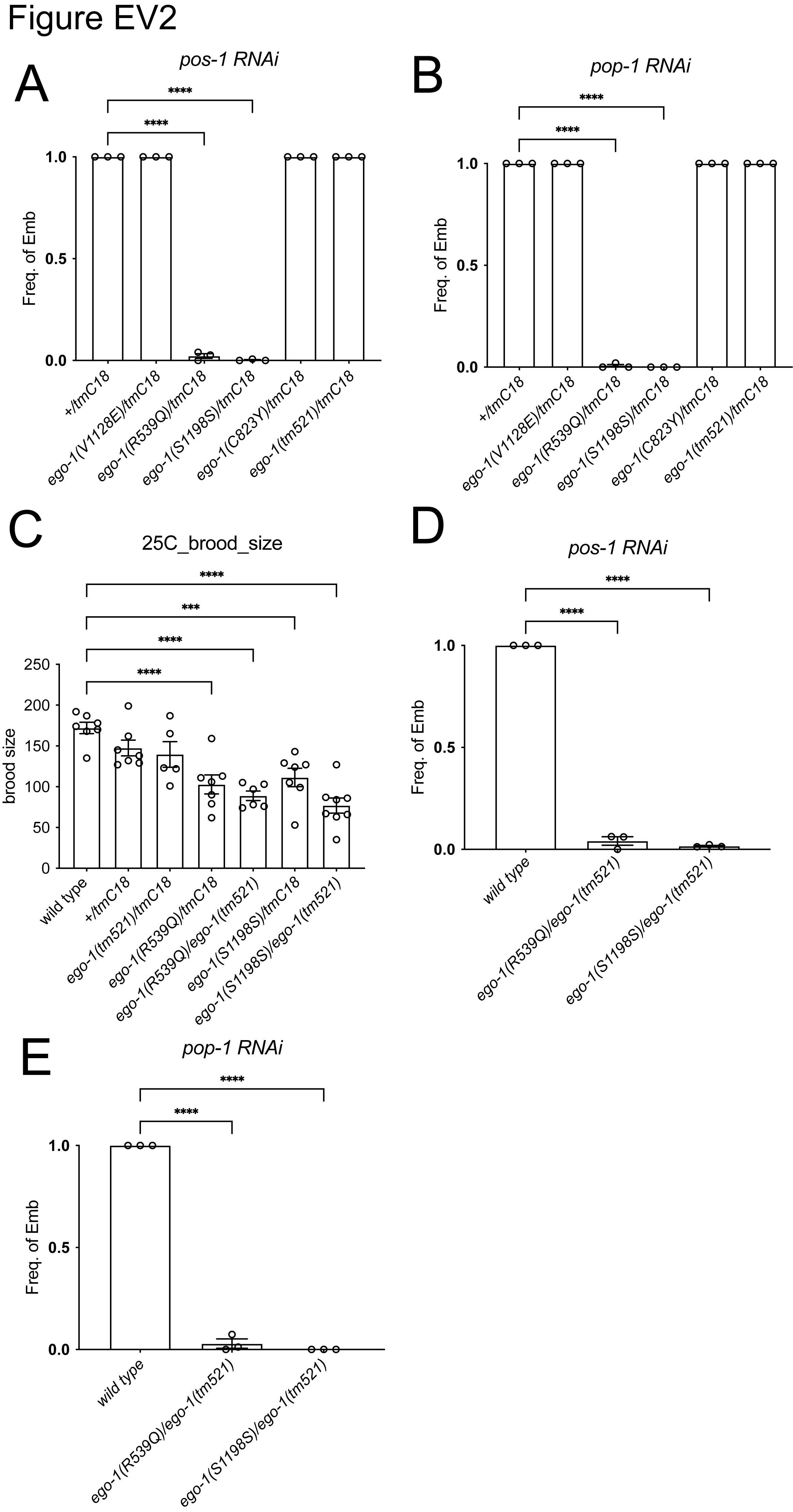
